# Unravelling the distribution of vectors of major vector-borne diseases in Koshi Province of Nepal: A concern of expansion in diverse geo-ecological and climatic regions

**DOI:** 10.1101/2025.06.02.657347

**Authors:** Lalita Roy, Surendra Uranw, Raja Ram Pote Shrestha

**Affiliations:** Tropical and Infectious Disease Centre, B. P. Koirala Institute of Health Sciences, Dharan, Nepal; Department of Internal Medicine, B. P. Koirala Institute of Health Sciences, Dharan, Nepal; World Health Organization, Country Office, Kathmandu, Nepal

## Abstract

**Background:** Vector-borne diseases, being major public health concerns, are either slated for elimination or projected for control in Nepal. One of the major challenges in controlling these VBDs is to halt their emergence and expansion in diverse geo-ecological and climatic regions. In this study, we collected vectors of major VBDs to understand their distribution, diversity, and association with ecological variances.

**Methodology/Principal findings:** A descriptive cross-sectional survey was conducted to collect the vectors in five districts situated at altitudes of up to 98 – 1,274 meters in Koshi Province, Eastern Nepal, representing three distinct geo-ecological and climatic regions: mountain, hills and lowlands during May and June 2023. Adult vectors were captured using CDC miniature light traps, BG sentinel traps, and manual aspirators. Association of vectors’ abundance in function of the geo-ecological and climatic variables was assessed in two vector species (*Culex quinquefasciatus* and *Phlebotomus argentipes*). We found the malaria vector *Anopheles annularis*, the lymphatic filariasis vector *Cu. quinquefasciatus* and visceral leishmaniasis vector *Ph. argentipes* from all three geo-ecological regions. The other vectors of the malaria parasite, *An. pseudowillmori* and *An. willmori*, and Japanese encephalitis vector *Cu. tritaeniorhynchus* were recorded from hilly districts only. Mean temperature and rainfall had positive effect on *Cu. quinquefasciatus* density and deleterious on *Ph. argentipes. Culex quinquefasciatus* and *Ph. argentipes* were captured in high abundance per household in hills (IRR = 1.23 and IRR = 13.00, respectively) and mountain (IRR = 1.96 and IRR = 4.00, respectively) as compared to lowland.

**Conclusion:** Two major vectors, *Cu. quinquefasciatus* and *Ph. argentipes* were indiscriminately present in all geo-ecological regions. Climatic variables favour vectors’ survival, distribution and growth in diverse altitudes, including areas previously impervious to the vectors and VBDs. Our findings alert the VBD control program and suggest regular monitoring, strengthening the existing surveillance system and evidence-based planning and implementation of vector control interventions in wider geo-ecological regions to prevent the disease transmission.

**Authors Summary:** The Government of Nepal is committed to eliminate and control major vector-borne diseases: malaria, visceral leishmaniasis, lymphatic filariasis, Japanese encephalitis and dengue as a public health problem by 2030. These diseases, once endemic to lowlands with tropical and subtropical climates only, have been spread to a wide geography, including hills and mountains. These high-altitude areas were considered unsuitable for the existence of vectors and transmission of pathogens. Considering the impact of climate change on founding a favorable environment for the establishment of viable populations of vectors for pathogens transmission and thus threatening all the efforts of VBDs control and elimination, we conducted a cross-sectional survey to explore the diversity of vectors in varied geo-ecological regions. We collected vectors from five districts situated at different altitudes and having fairly diverse geography. Major vectors of responsible of transmission of pathogens causing VL and LF were abundantly present in all geo-ecological regions, while vectors of malarial parasites and dengue virus were sparsely present in fewer areas. This study also examined the feasibility of the integrated vector survey in favour of maximising the available resources for entomological investigations. Further, this study provides updated information on the diversity of the vectors in the areas having cases of VBDs and suggests authorities for regular monitoring, strengthening the existing surveillance system and evidence-based planning and implementation of vector control interventions to prevent the disease transmission in wider geo-ecological regions.

## Background

Nepal is endemic to at least six vector-borne diseases; malaria, visceral leishmaniasis (VL), lymphatic filariasis (LF), Japanese encephalitis (JE), dengue and scrub typhus [1]. These diseases are prevalent in the poorest and most marginalized populations without receiving adequate resources for good management. Historically, VBDs were endemic to lowland, the geo-ecological region situated below 600 m asl and the area mostly covered with forest until early 20^th^ century. In recent years, VBDs are spread to other geo-ecological regions situated at higher altitudes: hills and mountains. These regions were once considered unsuitable to thrive the vector population and thus transmission of the pathogen was not anticipated. The geographical shift in distribution of VBDs is attributed to several factors, of which climate change is assumed to be the most important one. Changes in temperature, humidity and precipitation suited well for vector survival, maintaining the transmission cycle, and thus leading to a wider geographical spread of the pathogens, vectors and vector-borne diseases [2-5]. The situation ultimately imposes the threat of large-scale epidemics in relatively naïve susceptible populations, with overburdening of unprepared health systems for VBDs control and low-quality health care as a consequence [4, 6, 7].

Over the past decades, Nepal has been experiencing noticeable climate changes especially in two crucial climatic variables: temperature and precipitation. There is an observed increase in temperature with 1.5°C over the last two and a half decades in Nepal, as compared to 0.6°C at the global level [8]. Similarly, precipitation has increased significantly with 5.3% per decade over the last six decades, with a more rapid increase since the mid-1980s [9]. Recent climatic data demonstrated increase in rainfall with altitude on the windward side and decrease on the downwind side in the middle mountains. Analysis of rainfall shows average annual rainfall is 1883.8 mm in lowlands (below 1000m elevation) and 1959.6 mm in highlands (above 1000m elevation). The review suggested that July is the highest rain month (pre-monsoon to monsoon) and November (post monsoon) is the lowest [10].

Though, there is substantial information available on the reported of VBDs at different climatic and ecological regions of Nepal, detail investigation on the occurrence and bionomics of vectors of VBDs is still lacking. Inadequate vector surveillance data is a major limitation in the proper planning and implementation of the vector control programme in Nepal. Integrated vector monitoring or surveillance is one step closer to the planning and execution of integrated vector management (IVM), aiding economic benefit to vector control programmes for major VBDs [11].

In this survey, we collected the baseline information on vectors of major VBDs particularly malaria, VL, LF, JE and dengue in different geo-ecological and climatic variances in selected districts of Koshi Province, situated in eastern Nepal. Abundance of vector and non-vector species, association with the geo-ecological (mountain, hills, lowland and housing conditions) and climatic factors (temperature, rainfall and relative humidity) and spatial relationship between respective vector abundances and VBDs in the identified areas were also assessed.

## Methods

### Survey sites

Nepal topographically divided from north to south into three ecological regions: the high mountains, the hills and the lowland also known as “terai” (Fig 1). This topography is reflected by geographical and socio-cultural diversity as well. The country is administratively divided into 7 provinces and 77 districts. Though Nepal is well known for its mountains, over 80% of the total population of 30 million lives in lowland and harbour most tropical and subtropical diseases. We reviewed the epidemiological surveillance data of the Epidemiology and Disease Control Division (EDCD), Ministry of Health & Population, Government of Nepal and the VBDs patient database of the Koshi Province for the period 2020 – 2022. All VBDs cases from the last three years who named from different districts of Koshi province as their residence at the time of admission were listed and the five districts were selected for vector survey. These five districts are representative of the three geo-ecological regions; Morang and Sunsari in the lowland, Udayapur in the low-mid hill, Okhaldhunga in the high hill and Sankhuwasabha district in the mountainous region (Fig 1, Table 1). In each district, we selected two clusters/ villages based on the criteria: (i) reported of at least one of the major VBDs cases since 2020, (ii) accessibility on foot, and (iii) willing to support by local health authorities and community. The selected locations of vector survey districts and clusters from five districts is given in table 1.

**Fig 1.**
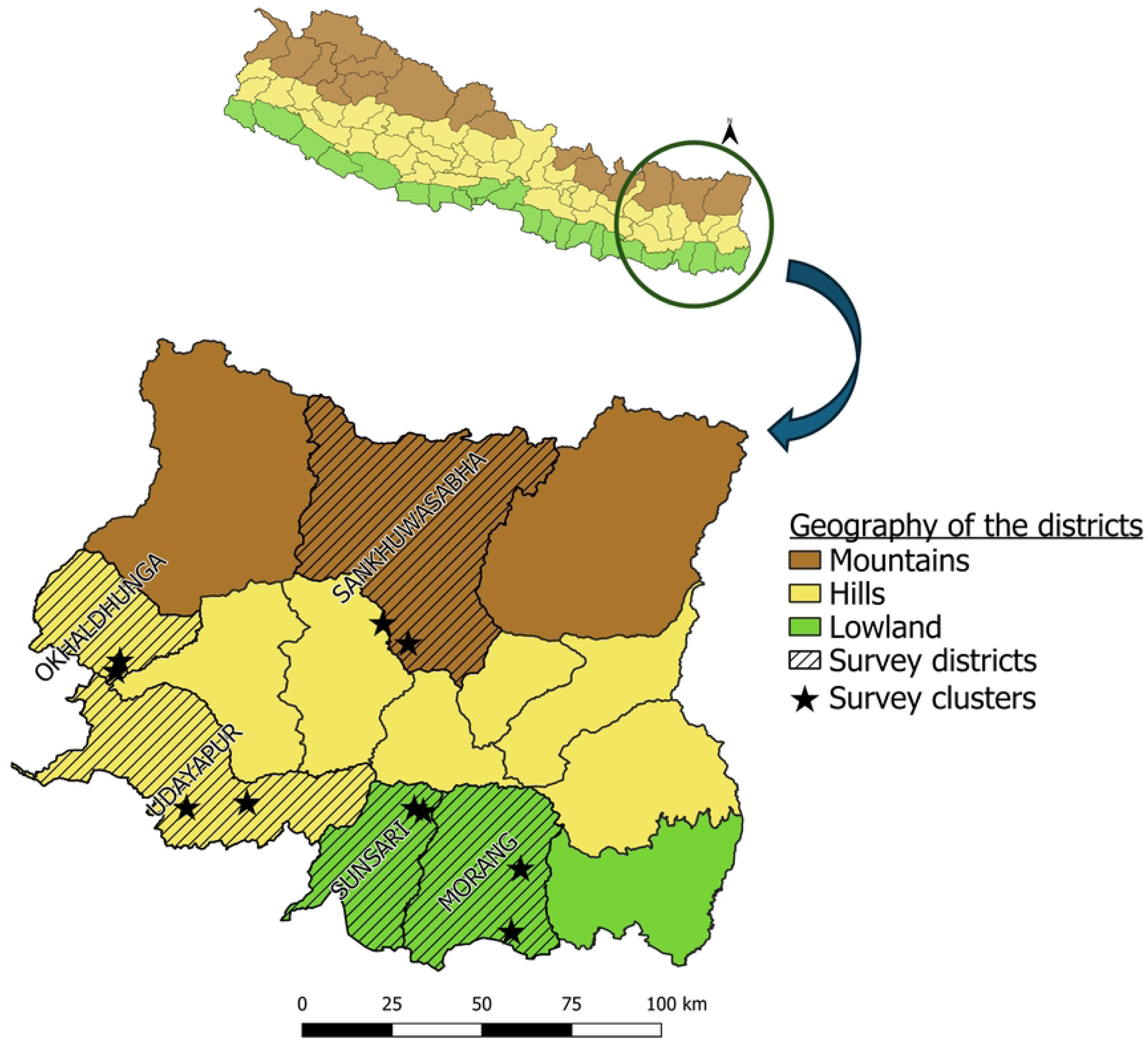
Location of vector survey districts and clusters in Koshi Province, Nepal, 2023.

**Table 1.**
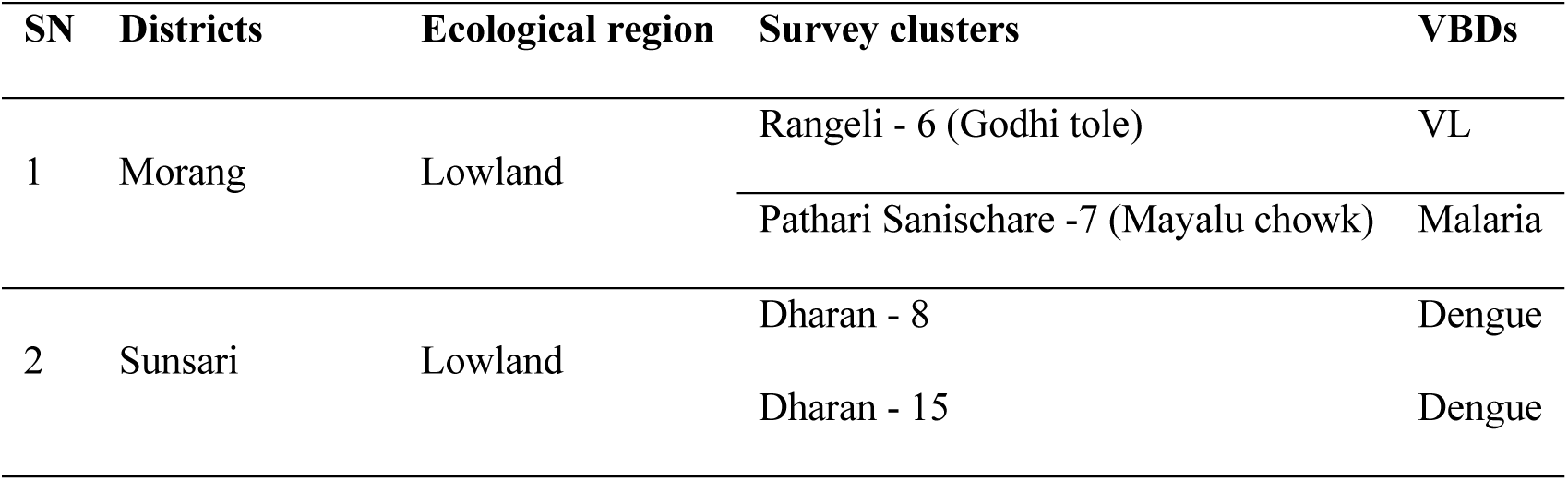
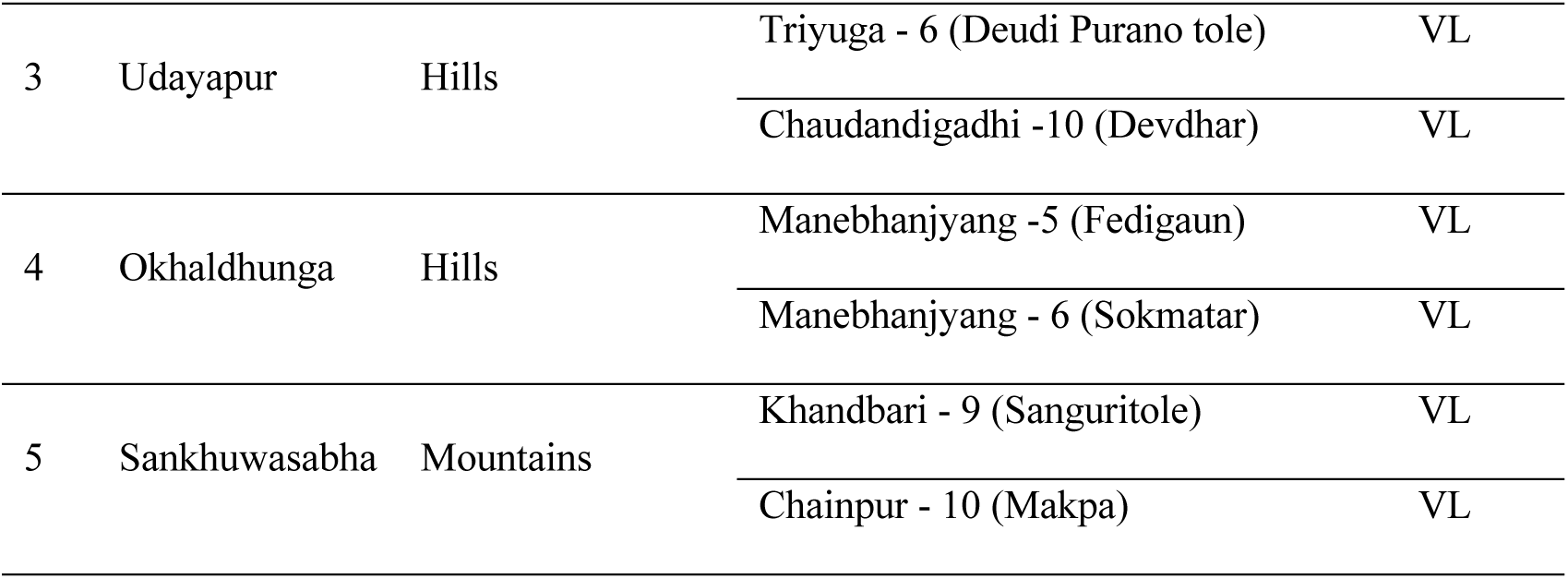
Details of the survey districts/clusters, Koshi Province, Nepal.

### Field survey

#### Collection of vectors

A descriptive cross-sectional entomology survey was conducted in the month of May and June 2023, timing purposively coincided with the first annual peak of vector density in lowland in Nepal. In each cluster, ten household structures (human dwelling, mixed dwelling, and cattle sheds) and nearby areas were explored for two consecutive days. Adult and immature stages of vector and other non-vector species were collected through various methods described below.

i. Adult *Anopheles* and *Culex* mosquitoes (vectors of the malarial parasite, LF, and JE) were collected both from indoor and outdoor human dwellings, cattle sheds, and mixed dwellings (where human beings and animals share the same roof in a structure) by using mouth aspirators. A cattle-baited trap (CBT) was used for outdoor mosquito collection at outside the household and/or nearby vegetation areas. Centre for Disease Control and Prevention (CDC) light traps were used for indoor mosquito collection along with sand flies.
ii. *Aedes* mosquitoes (vectors of dengue, chikungunya and zika virus): Five Biogents (BG) sentinel traps were placed outdoors near the households and the vegetation for the collection of *Aedes* mosquitoes.
iii. Sand flies (vectors of kala-azar): CDC light traps were placed indoors in human dwellings and mixed dwellings for sand fly and indoor mosquito collection. Resting sand flies were collected by mouth aspirators in the early morning during the survey. In each selected cluster, 10 CDC light traps (LTs) were fixed in 10 selected households, one in each house or cattle shed, for indoor mosquito and sand fly collections. Light traps were installed and operated from 18 hrs the evening to 6 hrs the next morning. The same process was repeated for the next day to have two consecutive nights of collection. Additionally, the resting sand flies were collected for two consecutive mornings by mouth aspiration for 15 minutes in the same households and the cattle sheds where LTs were installed. Mosquitoes and sand flies were collected and dry preserved in tubes with silica gel, labeled with cluster code, site and collection method. The tubes were transported to the entomology laboratory at B.P. Koirala Institute of Health Sciences, Dharan, Nepal. Mosquitoes and sand flies were identified up to species-level using regional keys [12-20], a stereoscope and a light microscope. After identification, mosquitoes were dry preserved in tubes with silica gel, and sand flies in tubes with 80% ethanol at species and sex levels for each cluster.

#### Immature stage survey

A detailed survey targeting the immature stages of *Aedes* mosquitoes were conducted in two wards of Dharan sub-metropolitan city ward number 8 and 15 in Sunsari, where increasing number of dengue fever cases were recorded in recent months during the survey. Larval and pupal sampling methods were adapted from the standard operating procedure (SOP) and guidelines developed by WHO [21, 22]. Random houses in these wards and public places were searched indoors and outdoors for water-holding containers and the presence of mosquitoes’ immature stages. Overhead tanks were not searched due to inconvenience and safety reasons. Besides the water-holding containers for household purposes, the discarded plastic and metal containers, tyres, flower vases, plates kept under flower pots, kitchen gardens, mud pots, gallons, tree holes wherever possible, and any form of utensils that can hold water were also searched for the presence of immature stages. Positive containers were sampled and immature stages (larvae and pupae) were collected and transported to the entomology laboratory for further rearing to adult stage and then identified to species level.

#### Housing characteristics and geo-ecological information

All survey households were geo-referenced by a Global Positioning System (GPS) device, specifying longitude, latitude and altitude of the houses. Trained entomology technicians interviewed each head of household to collect the details of housing structures, presence of cattle and other domestic animals, surrounding vegetation, water bodies, etc. for the household where CDC LTs were kept. A semi-structured questionnaire was used to record the information on feeding and resting sites of mosquitoes and sand flies collected.

#### Climate data

Climatic data such as daily records of rainfall, relative humidity (%), minimum and maximum temperature (°C) for the period of one year (July 2022 – June 2023) for each survey cluster were obtained from the nearest meteorological stations of the Department of Hydrology and Meteorology, Government of Nepal. The meteorological stations were located nearby the survey villages/clusters (<10 kilometers) in the lowland and less than 5 kilometers away in upland (hills).

#### Data management and analysis

All data collected in the field was entered in databases made in Epi Info version 3.5.1. Descriptive analysis was performed for household characteristics and climatic data (temperature, rainfall and relative humidity). Abundance and species richness (S) of vector and non-vector species were represented by absolute numbers. Species diversity and dominance or uniformity in distribution of the species at the district level were represented in Shannon-Wiener diversity index (H’) and Pielou’s evenness index (J) [23, 24]. H’ and J are calculated using function ‘diversity’ from a package “vegan” in R [25]. Mathematical calculation was done using the following indices:

a. Shannon-Wiener diversity index (H’) = -Σpi * ln(p_i_)

Where, Σ: Sum, ln: Natural log and pi = n_i_/N (n_i_ = the number of individuals of a species and N = Total number of individuals)

b. Pielou’s evenness index (J) = H’/ln(S)

Where, H’ = Shannon-Wiener diversity index and S is the total number of species in a sample

Interpretation of Shannon-Wiener diversity index (H’) was done as the higher the value of H’, the higher the diversity of species and lower the value, the lower the diversity and if value of H’ is 0 then only one species is present in that community. Pielou’s evenness index (J) ranges from 0 to 1. Higher the value of J, higher the level of evenness in the abundance of different species present in a particular community while lower value represents either one or only few species are present in abundance. A landscape map was prepared to illustrate the relative abundances (proportion) of the vector species according to the survey districts and elevation. *Stegomyia* indices (HH index, container index, Breteau index and pupae per person) were calculated for the immature stages of *Aedes* mosquitos collected from urban area of Dharan sub-metropolitan in Sunsari.

As the mosquitoes and sand fly count at house hold level was over-dispersed and had shown non-normal distribution, i.e., variance was larger than the mean value, we fitted generalized linear models (GLM) with a negative binomial distribution to assess the association of the vector abundance in function of the explanatory variables like ecological regions, method of collection, collection sites, climatic variables, household structures, and surrounding ecological features. Spearman’s correlation was assessed between each vector species with the mean temperature (°C), mean relative humidity (%) and cumulative rainfall (mm) of the preceding one month and the month when the survey was conducted, i.e., April and May 2023, before incorporating them in the model. Final model was fitted separately for each vector species. The vector species, *An. annularis, An. pseudowillmori, An. willmori, Cu. tritaeniorhynchus, Ae. aegypti* and *Ae. albopictus* were not included in the final model due to their low number of collections. Hence, the model was fitted with two vector species with plausible collection; *Cu. quinquefasciatus* and *Ph. argentipes*. The calculation was done using the function ‘glm.nb’ from a R package “MASS” [26]. Results of the analysis are interpreted as an incidence rate ratio (IRR) and confidence interval (CI) at 95%.

The incidence rate and vector abundance gradient map for LF and VL were constructed using QGIS version 3.36. The disease incidence rates for LF and VL were calculated per 10,000 population at the district level from the available data and the national surveillance data collected in 2022 and 2023. The association of the vector abundance and the presence of VBDs at the district level were analyzed with the same GLM method explained above and the outcome has explained in terms of IRR and confidence interval at 95%.

#### Ethical consideration

Ethical approval to conduct this study was obtained from the Ethical Review Board of the Nepal Health Research Council (NHRC), Kathmandu, Nepal (268/2022P) and the ethical review committee of the WHO South East Asia Regional Office, New Delhi, India.

## Results

### Characteristics of districts, households and surroundings in the survey clusters

The survey districts, namely Sunsari, Morang, Udaypur, Sankhuwasabha and Okhaldhunga are endemic to visceral leishmaniasis, malaria and lymphatic filariasis. We retrieved cumulative 101 past VL cases in the document review, five from Sunsari and Sankhuwasabha, 11 from Morang, 12 from Udaypur and 68 from Okhaldhunga during the period of 2020-2022. Only one malaria case was reported from Morang in 2020. Lymphatic filariasis cases were reported from all five districts, but they were not from the survey clusters. Dengue fever is an emerging threat in all these clusters.

Elevation of survey clusters ranged from 98 m asl in one of the cluster in the lowland (Morang) to 1,274 m asl in a cluster situated in high hill (Okhaldhunga). The key characteristics of survey districts and the households where light traps were kept, and their surrounding areas, are shown in Table 2.

**Table 2.** Key characteristics of the survey districts and households in Koshi Province, 2023.

| General characteristics | Morang | Sunsari | Udayapur | Okhaldhunga | Sankhuwasabha |
| --- | --- | --- | --- | --- | --- |
| Ecological region | Lowland | Lowland | Low-mid hills | High hills | Mountain |
| Urbanization of clusters | Rural | Urban | Rural | Rural | Rural |
| Altitude of survey clusters (m asl) (min-max) | 98-139 | 309-323 | 437-623 | 1155-1274 | 832-1011 |
| Endemic for VBDs* | Yes | Yes | Yes | Yes | Yes |
| **Household** | **N = 20 (%)** | **N = 20 (%)** | **N = 20 (%)** | **N = 20 (%)** | **N = 20 (%)** |

| **characteristics** |  |  |  |  |  |
| --- | --- | --- | --- | --- | --- |
| Type of roof |  |  |  |  |  |
| Cement | 3 (15%) | 0 (0%) | 0 (0%) | 0 (0%) | 0 (0%) |
| Thatch (Straw and bamboo) | 0 (0%) | 2 (10%) | 1 (5%) | 6 (30%) | 9 (45%) |
| Tiles | 0 (0%) | 0 (0%) | 1 (5%) | 0 (0%) | 0 (0%) |
| Tin | 17 (85%) | 18 (90%) | 18 (90%) | 14 (70%) | 11 (55%) |
| Type of wall |  |  |  |  |  |
| Cemented | 8 (40%) | 6 (30%) | 4 (20%) | 5 (25%) | 1 (5%) |
| Unplastered brick | 1 (5%) | 2 (10%) | 1 (5%) | 2 (10%) | 0 (0%) |
| Mud | 11 (55%) | 10 (50%) | 15 (75%) | 13 (65%) | 19 (95%) |
| Tin | 0 (0%) | 2 (10%) | 0 (0%) | 0 (0%) | 0 (0%) |
| Type of Floor |  |  |  |  |  |
| Cement | 9 (45%) | 10 (50%) | 6 (30%) | 5 (25%) | 3 (15%) |
| Earthen | 11 (55%) | 10 (50%) | 14 (70%) | 15 (75%) | 17 (85%) |
| Presence of ventilation | 3 (15%) | 7 (35%) | 2 (10%) | 4 (20%) | 1 (5%) |
| Presence of cattle | 7 (35%) | 0 (0%) | 10 (50%) | 12 (60%) | 11 (55%) |
| Presence of cow dung | 7 (35%) | 3 (15%) | 12 (60%) | 15 (75%) | 14 (70%) |
| Presence of goats | 9 ((45%) | 5 (25%) | 9 (45%) | 18 (90%) | 14 (70%) |
| Presence of pigs | 7 (35%) | 2 (10%) | 5 (25%) | 4 (20%) | 12 (60%) |
| Presence of agricultural field | 18 (90%) | 2 (10%) | 18 (90%) | 10 (50%) | 18 (90%) |
| Presence of vegetable field | 14 (70%) | 7 (35%) | 15 (75%) | 14 (70%) | 14 (70%) |
| Presence of mixed orchard | 20 (100%) | 13 (65%) | 16 (80%) | 18 (90%) | 19 (95%) |
| Presence of river nearby | 10 (50%) | 14 (70%) | 18 (90%) | 14 (70%) | 6 (30%) |
| Presence of pond nearby | 10 (50%) | 1 (5%) | 9 (45%) | 6 (30%) | 0 (0%) |
| **Mean number of domestic animals (± sd) per household** |  |  |  |  |  |
| Cattle | 1.25 ($\pm 1.97$) | 0.05 ($\pm 0.22$) | 2.3 ($\pm 3.13$) | 2.5 ($\pm 2.40$) | 2.35 ($\pm 2.41$) |
| Goats | 2.05 ($\pm 3.3$) | 1.85 ($\pm 4$) | 1.95 ($\pm 3.89$) | 13.5 ($\pm 12.9$) | 4.2 ($\pm 5.32$) |
| Pigs | 1.15 ($\pm 2.23$) | 0.15 ($\pm 0.49$) | 0.5 ($\pm 1.1$) | 0.25 ($\pm 0.55$) | 1.25 ($\pm 1.21$) |

*\*visceral leishmaniasis, malaria, lymphatic filariasis, dengue fever*

### Status of climatic variables

#### Temperature

The average daily maximum temperature of the surveyed clusters in three different ecological regions varied from 31°C in lowland (Sunsari) to 23°C in high hills (Okhaldhunga). The average daily maximum temperature was reported approximately 35°C in the month of June in lowland (Morang) compared to 28°C in high hills (Okhaldhunga). In the same month, it was observed that there was a temperature variation on average 7°C daily between lowland and highlands or hills. The average daily minimum temperature experienced in lowland (Rangeli, one of the cluster in Morang) was about 9°C in January compared to 7°C in high hills (Okhaldhunga) in the same month (S1 Fig).

#### Relative humidity

Average daily relative humidity varied from 55.7% in April to 89.1% in September in the survey clusters/districts. April was marked as the driest month and September as the wettest month in terms of moisture present in air. The observed relative humidity varied from 70.2% in lowland (Morang) to 80.8% in high hills (Okhaldhunga). Average annual relative humidity was found to be higher in high hills (Okhaldhunga) compared to all other surveyed districts (S2 Fig).

#### Rainfall

Annual rainfall also varied according to the ecological regions; the lowest annual rainfall (1451.6mm) was observed in one of the survey cluster located in the lowland (Rangeli, Morang) and highest rainfall (2037.7mm) in the mid hills (Udayapur). Higher average daily rainfall was observed in June to September (S3 Fig).

### Entomological findings

#### Abundance and types of vector species

The total number of mosquitoes and sand flies captured was 3,867, of which, vector species comprised 77.4% (n = 2,994); morphologically identified and segregated into six genera with 28 species of mosquitoes and two genera with three known species of sand flies. Amongst the captured vector species, few specimens of genus *Phlebotomus* (n=11) could not be identified up to the species level. Variation in the species composition of mosquitoes was evident in surveyed clusters and districts. Diversity index was higher in Okhaldhunga for all species collected (H’ = 1.45) and also for the vector species (H’ = 0.57). Species richness for both vector and non-vector species was higher in Udayapur (S = 20). For vector species only, species richness was higher in Okhaldhunga (S = 5). Pielou’s evenness index illustrated the fact that one or few species were dominant, and others were present with nominal density at the time of collection (Tables 3 and 4).

**Table 3.** Diversity index, evenness index, species richness and abundance of all vector and non-vector species in five districts of Koshi Province, 2023.

| Districts | Both vector and non-vector species |  |  |  | Vector species only |  |  |  |
| --- | --- | --- | --- | --- | --- | --- | --- | --- |
|  | H' | J | S | A | H' | J | S | A |
| Morang | 0.96 | 0.36 | 14 | 906 | 0.10 | 0.09 | 3 | 704 |
| Sunsari | 1.23 | 0.49 | 12 | 352 | 0.51 | 0.37 | 4 | 163 |
| Udayapur | 0.90 | 0.30 | 20 | 1259 | 0.09 | 0.07 | 4 | 1021 |
| Okhaldhunga | 1.45 | 0.55 | 14 | 395 | 0.57 | 0.36 | 5 | 254 |
| Sankhuwasabha | 0.70 | 0.23 | 20 | 955 | 0.20 | 0.14 | 4 | 852 |

*Note: $H'$ – Shannon's diversity index, $J$ – Pielou's species richness index, $S$ – Species richness, $A$ - Abundance*

**Table 4.** Distribution and abundance of vector and non-vector species in 10 clusters of five survey districts in Koshi province, 2023.

| Districts | Morang |  | Sunsari |  | Udayapur |  | Okhaldhunga |  | Sankhuwasabha |  | Total,<br>n (%) |
| --- | --- | --- | --- | --- | --- | --- | --- | --- | --- | --- | --- |
| Clusters | Pathari<br>Sanischare-7 | Rangeli-6 | Dharan-8 | Dharan-15 | Chaudandigadhi-10 | Triyuga-6 | Manebhanjyang-5 | Manebhanjyang-6 | Chainpur-10 | Khandbari-9 |  |
| Ecological region | Lowland | Lowland | Lowland | Lowland | Hills | Hills | Hills | Hills | Mountain | Mountain |  |
| Species |  |  |  |  |  |  |  |  |  |  |  |
| *Aedes aegypti* * | - | - | 5 | 7 | - | - | - | - | - | - | 12 (0.31) |
| *Aedes albopictus* * | - | - | 1 | - | - | - | - | - | - | 1 | 2 (0.05) |
| *Aedes pseudotaeniatus* | - | - | - | - | - | - | 1 | - | - | - | 1 (0.03) |
| *Aedes sp.* | - | - | - | - | - | - | - | 1 | - | - | 1 (0.03) |
| *Anopheles aconitus* | - | - | - | - | - | 1 | - | - | - | - | 1 (0.03) |
| *Anopheles annularis* * | 4 | 1 | - | - | - | 9 | 1 | - | 1 | 2 | 18 (0.47) |
| *Anopheles barbirostris* | - | - | - | - | - | 2 | - | - | - | - | 2 (0.05) |
| *Anopheles culicifacies* | 2 | 2 | 2 | 1 | 22 | 62 | 1 | 1 | 1 | 4 | 98 (2.53) |
| *Anopheles pseudowillmori* * | - | - | - | - | - | - | 12 | 2 | - | - | 14 (0.36) |
| *Anopheles stephensi* | - | - | 5 | 1 | - | 1 | - | - | - | - | 7 (0.18) |
| *Anopheles subpictus* | 3 | 66 | 1 | - | - | 5 | - | - | 3 | - | 78 (2.02) |
| *Anopheles* UN1 | - | - | - | - | - | 1 | - | - | - |  | 1 (0.03) |
| *Anopheles vagus* | 7 | 27 | - | - | 2 | 29 | - | - | - | 3 | 68 (1.76) |
| *Anopheles willmori* * | - | - | - | - | - | - | 5 | - | - | - | 5 (0.13) |
| *Armegeres kesseli* | 22 | - | - | - | 6 | 18 | - | - | - | 22 | 68 (1.76) |
| *Armegeres kuchingensis* | - | - | - | - | - | 1 | - | - | - | - | 1 (0.03) |
| *Armegeres subalbatus* | - | 2 | 3 | 3 | 1 | - | - | - | - | 4 | 13 (0.34) |
| *Culex bitaeniorhynchus* | 1 | - | - | - | - | 3 | - | - | - | - | 4 (0.10) |
| *Culex fuscocephala* | 1 | - | 3 | 1 | - | 16 | 1 | - | 5 | 27 | 54 (1.40) |
| *Culex infula* | 1 | - | - | - | - | - | - | - | - | - | 1 (0.03) |
| *Culex mimulus* | - | - | - | - | - | 1 | - | - | - | - | 1 (0.03) |
| *Culex quinquefasciatus* * | 256 | 435 | 66 | 75 | 3 | 1003 | 7 | 10 | 96 | 718 | 2669 (69.02) |
| *Culex sinensis* | 1 | - | - | - | - | - | - | - | - | - | 1 (0.03) |
| *Culex tritaeniorhynchus* * | - | - | - | - | - | 2 | - | - | - | - | 2 (0.05) |
| *Culex vagans* | - | - | - | 1 | - | - | - | - | - | - | 1 (0.03) |
| *Culex whitei* | - | 2 | - | - | - | 2 | - | - | - | - | 4 (0.10) |
| *Mansonia uniformis* | - | - | - | 1 | - | 2 | - | - | - | - | 3 (0.08) |
| *Phlebotomus* (Adlerius)<br>spp. | - | - | - | - | - | - | 2 | 1 | - | - | 3 (0.08) |
| *Phlebotomus argentipes* * | 2 | 6 | 4 | 5 | 2 | 2 | 43 | 174 | 27 | 7 | 272 (7.03) |
| *Phlebotomus major* s.l. | - | - | - | - | - | - | 29 | 20 | 3 | 1 | 53 (1.37) |
| *Phlebotomus* spp. | - | - | - | - | - | - | 7 | 4 | - | - | 11 (0.28) |
| *Sergentomyia* spp. | 42 | 23 | 14 | 153 | 23 | 40 | 18 | 54 | 1 | 29 | 397 (10.27) |
| *Uranotaenia* sp. complex |  |  |  |  |  |  | 1 |  |  |  | 1 (0.03) |
| **Grand Total** | **342** | **564** | **104** | **248** | **59** | **1200** | **128** | **267** | **137** | **818** | **3867** |

Note: \*Vector species

#### Distribution of vector species among the districts and altitudinal gradient

The known malaria vectors in Nepal, *An. annularis* were captured from all the surveyed clusters/districts except Sunsari situated at 98m to >1,000m asl (lowlands to highlands). The other vectors for the malaria parasite, *An. pseudowillmori* and *An. willmori* were also recorded at 1,200m altitude in Okhaldhunga only. The vector transmitting *Wuchereria bancrofti* microfilariae causing lymphatic filariasis, *Cu. quinquefasciatus* is found at 98 to 1,274m asl during this survey. Similarly, the vector transmitting the virus causing Japanese encephalitis, *Cu. tritaeniorhynchus*, was found at 632m asl in Udayapur. Both vectors, *Ae. aegypti* and *Ae. albopictus* transmitting dengue virus were captured at 318m in Sunsari and only *Ae*. *albopictus* at 832m asl in Sankhuwasabha. *Phlebotomus argentipes*, the vector of *Leishmania donovani* parasites, was recorded at 98 to 1,274m asl. The vector sand fly abundance was four times higher in surveyed clusters of Okhaldhunga with an altitude >1000m asl. Other suspected sand fly vectors of *Leishmania* spp., *Ph*. *major* sensu lato and *Ph. (Adlerius)* sp. were also recorded from Okhaldhunga and Sankhuwasabha at altitudes from 832 to 1,011m asl. The details of the vector distribution in the survey districts are illustrated in landscape maps (Fig 2 and Fig 3).

**Fig 2.**
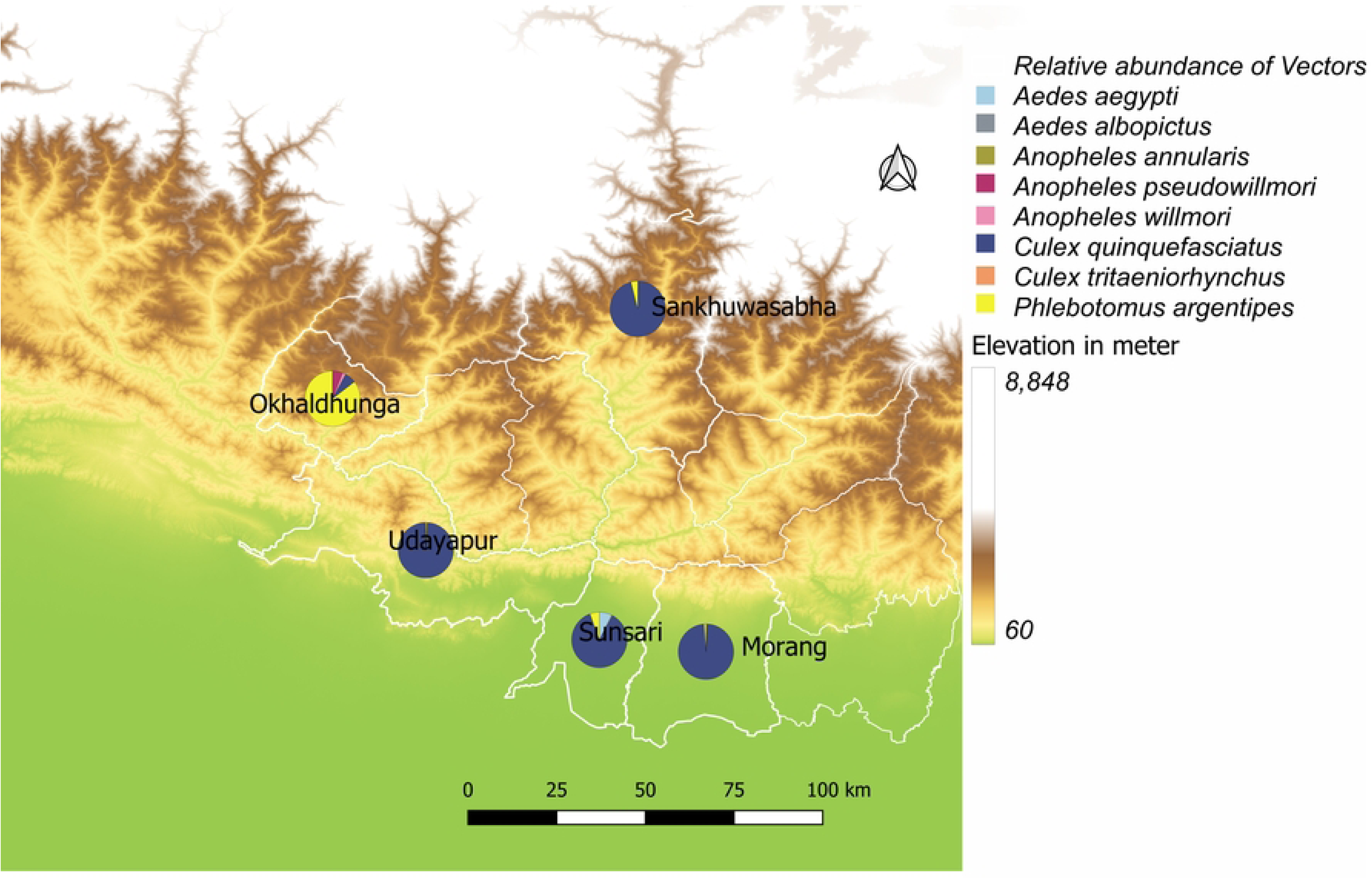
Location where relative abundance (proportion) of the vector species captured in survey districts (Map showing the elevation of the landscape; brown colour-high elevation and green colour-low elevation).

**Fig 3.**
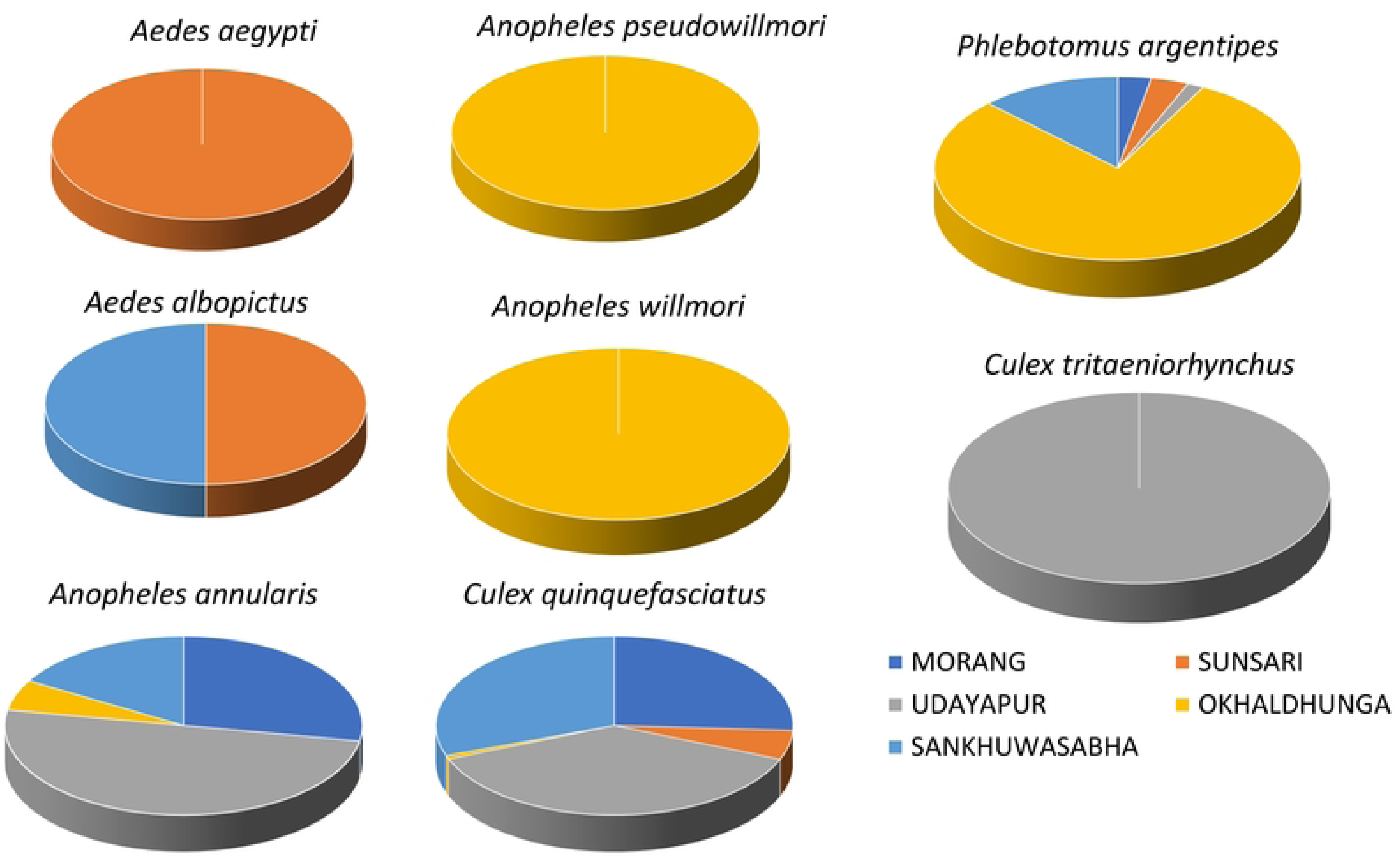
Relative distribution (proportion) of vector species in five survey districts, 2023.

#### Immature stages of *Aedes* mosquitoes

In two selected wards of Dharan sub-metropolitan city, the dengue outbreak was reported in 2019. In both wards, a total of 434 wet containers in 135 households (including few public places) with 525 inhabitants were inspected for the *Aedes* larvae and pupae. Altogether, 144 wet containers from 81 households were positive for the immature stages (Table 5). The household index (HI) was found to be 60% (81/135*100), the container index (CI) was 33.18% (144/343*100), the Breteau index (BI) was 106.67 (144/135*100). The pupae per person was 0.84 based on 443 pupae collected from the positive containers. These high *Stegomyia* indices (HI, CI, BI and PPP) were indicative of the outbreak situation for dengue fever in surveyed areas.

**Table 5.** Types of wet containers searched and their contribution to larval productivity.

| Types of wet containers | Searched containers (%) | Positive containers (%) |
| --- | --- | --- |
| Bowl | 6 (1.38) | 6 (4.17) |
| Bucket | 66 (15.21) | 11 (7.64) |
| Cement tank | 2 (0.46) | 2 (1.39) |
| Ceramic earthen jar | 5 (1.15) | 2 (1.39) |
| Coconut shell | 2 (0.46) | 0 (0.00) |
| Discarded bottles | 15 (3.46) | 5 (3.47) |
| Discarded plastic container | 15 (3.46) | 10 (6.94) |
| Discarded Tins | 6 (1.38) | 0 (0.00) |
| Ditch | 2 (0.46) | 1 (0.69) |
| Drum | 242 (55.76) | 63 (43.75) |
| Flower vase | 48 (11.06) | 27 (18.75) |
| Gallon | 2 (0.46) | 1 (0.69) |
| Mud pot | 2 (0.46) | 1 (0.69) |
| Plastic jar | 7 (1.61) | 5 (3.47) |
| Tyre | 14 (3.23) | 10 (6.94) |

#### Association of vector abundance with geo-ecological and climatic variables

Two vector species; *Cu. quinqfasciatus* (n = 2,669, 89.14% and *Ph. argentipes* (n = 272, 9.01%) were considered for the regression analysis as only they were present in plausible numbers for a valid interpretation as compared to other vector species (n = 53, 1.77%). Association of the geo-ecological and climatic variables are represented separately.

**For *Cu. quinquefasciatus*:** The effect of topography had a significant effect on the mean abundance of *Cu. quinquefasciatus*. Higher collections were recorded per household in hills (IRR = 1.23, CI at 95% = 0.53 – 2.87) and mountains (IRR = 1.96, CI at 95% = 0.73 – 5.91) as compared to the lowland. The result also indicated the existence of higher density of these vector in higher altitudes. CDC light trap was found to be excellent method of vector collection compared to aspirator (IRR = 0.05, CI at 95% = 0.03 – 0.08) and BG sentinel traps (IRR = 0.06, CI at 95% = 0.01 – 0.60). Vector density was found to be very low in cattle sheds, mixed dwellings and outdoors as compared to human dwellings (Table 6).

**Table 6.**
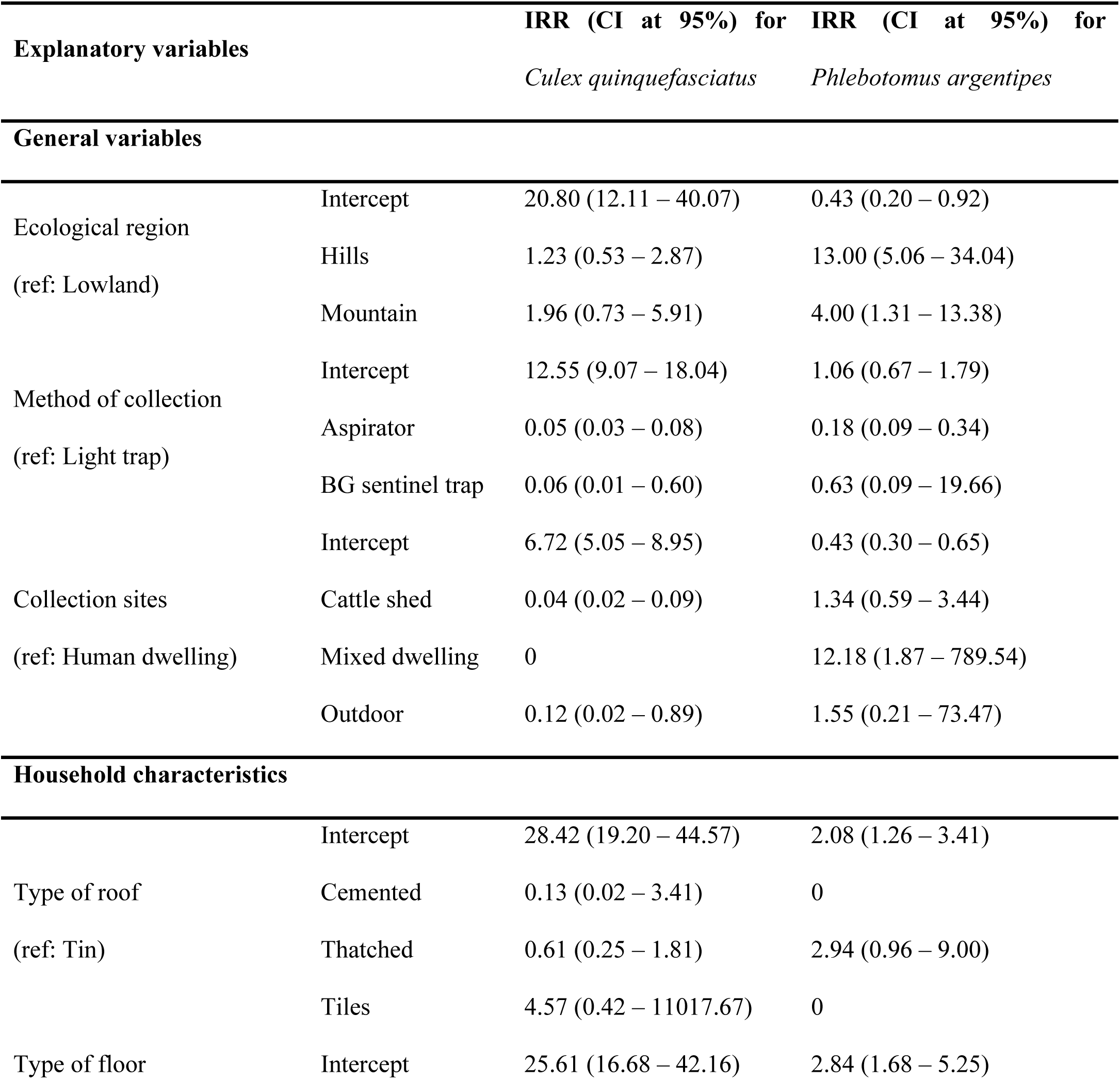
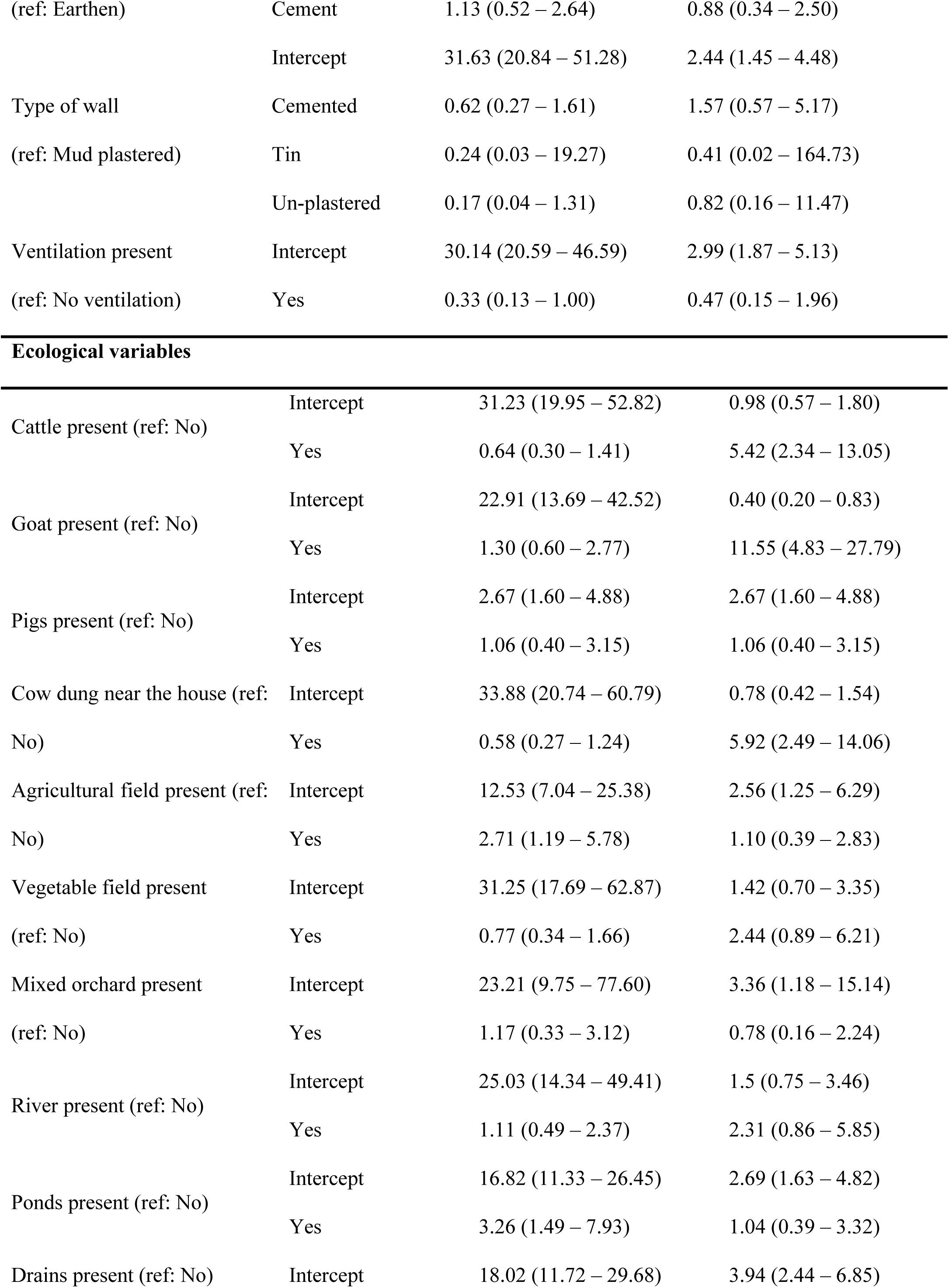
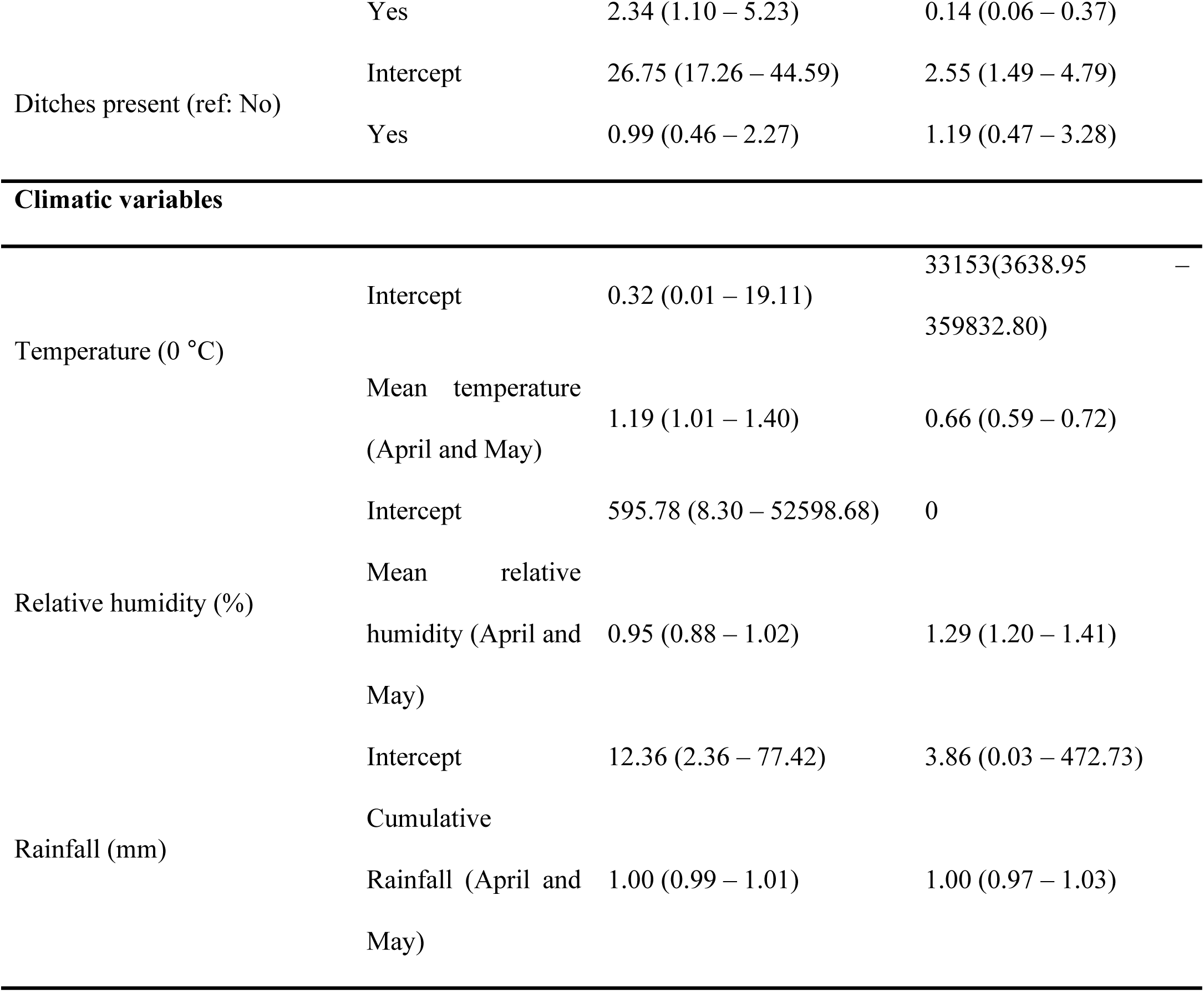
Association between geo-ecological and climatic factors with *Cu. quinquefasciatus* and *Ph. argentipes* density.

Considering the household structures, high vector density was recorded in the houses with tiled roofs, mud walls and cemented floor than other types of roof, wall or floor (Table 6). Vector density per household was found to be less in well ventilated rooms (IRR = 0.36, CI at 95% = 0.14 – 1.13) as compared to houses without proper ventilation. We observed the increasing effect *Cu. quinquefasciatus* density per household with the presence of goats, pigs, agricultural field, mixed orchard with variety of tropical and subtropical plants present nearby the household, presence of river, ponds and drains (Table 6).

**For *Ph. argentipes*:** A significant effect of topography has been observed with *Ph. argentipes* density which were collected almost 13 times higher in hilly and four times in mountainous districts as compared to lowlands. CDC light traps were found to be a highly efficient method of sand fly collection than mouth aspiration and BG sentinel traps. Mixed dwellings were highly productive in vector density (IRR = 12.18, CI at 95% = 1.87 – 789.54). Houses with thatched roofs, cemented walls, earthen floors, and poor or no ventilation showed an increasing effect on the vector density. Other ecological factors like the presence of cattle, goats, agricultural fields, vegetable fields, rivers, ponds, and ditches showed increasing effects on *Ph. argentipes* density per household (Table 6).

We also observed a weak but significant correlation with the *Cu. quinquefasciatus* density and mean temperature (r = 0.38, p<0.001) and rainfall (r = 0.11, p<0.001) while inversely proportional to the mean relative humidity recorded in the month of April and May (r = -0.32, p < 0.001). However, weak correlation was observed with rainfall (r = 0.11, p = 0.27). When these climatic factors were fitted into the model, the mean temperature recorded in the month of April and May had an increasing effect on the vector density (IRR = 1.19, CI at 95% = 1.01 – 1.40). Overall, the rainfall had negligible increasing effect on the vector density but while analysed at ecological region, the model demonstrated the increasing effect in hills and mountain but decreasing in the lowland. Another climatic variable, relative humidity had a decreasing effect on the density of these vectors per household in all ecological regions (Fig 4A, B and C).

**Fig 4.**
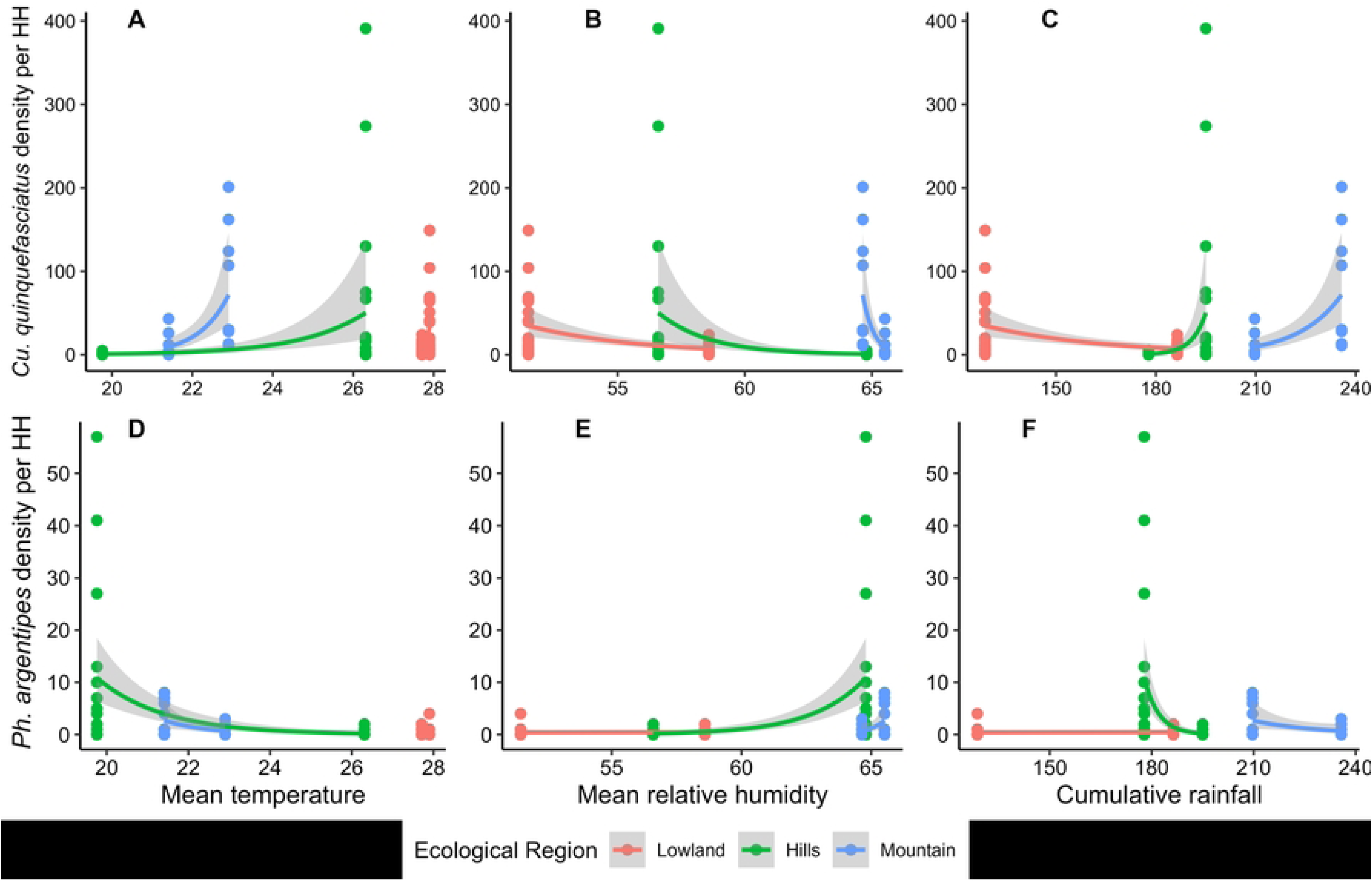
Scattered plots and regression lines showing effects of climatic variables on vector density in three ecological regions. Panels A, B and C show effects of climatic variables on *Cu. quinquefasciatus* density per HH (household) and panels D, E and F show effects on *Ph. argentipes* density per HH. Dots represent the data points, line represents the generalized regression line with negative binomial distribution and shaded area indicates the standard error of the regression line.

*Phlebotomus argentipes* density showed negative correlation with temperature (r = - 0.51, p<0.001), a weak but significant correlation with relative humidity (r = 0.50, p<0.001) and negative insignificant relation with rainfall (r = -0.10, p = 0.32). While fitted in the model with climatic data of April and May, the mean relative humidity showed an increasing effect (IRR = 1.29, CI at 95% = 1.20 – 1.41), the mean temperature had a decreasing effect and the cumulative rainfall showed no effect on the vector density (Table 6, Fig 4D, E and F).

#### Spatial relationship between vector abundance and vector-borne diseases

The high disease incidence for LF was well coincided with the high *Cu. quinquefasciatus* abundance in Udayapur (Fig 5A, B). The LF incidence rate was not available in the national line list for Morang and Sankhuwasabha. A similar pattern was also observed for VL; the district with the highest incidence rate had the highest number of *Ph. argentipes* (Fig 6A, B).

**Fig 5.**
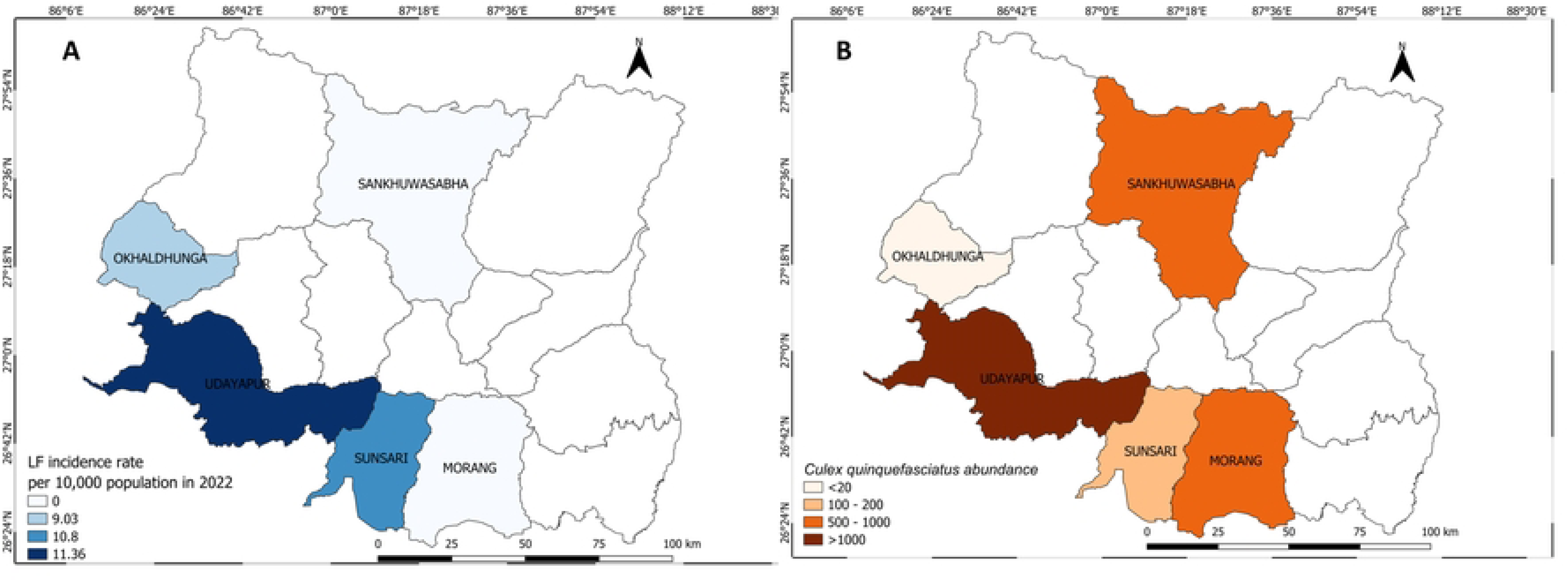
Lymphatic filariasis incidence rate in 2022 (A) and *Cu. Quinquefasciatus* abundance (B) during the survey in five study districts in Koshi province, 2023.

**Fig 6.**
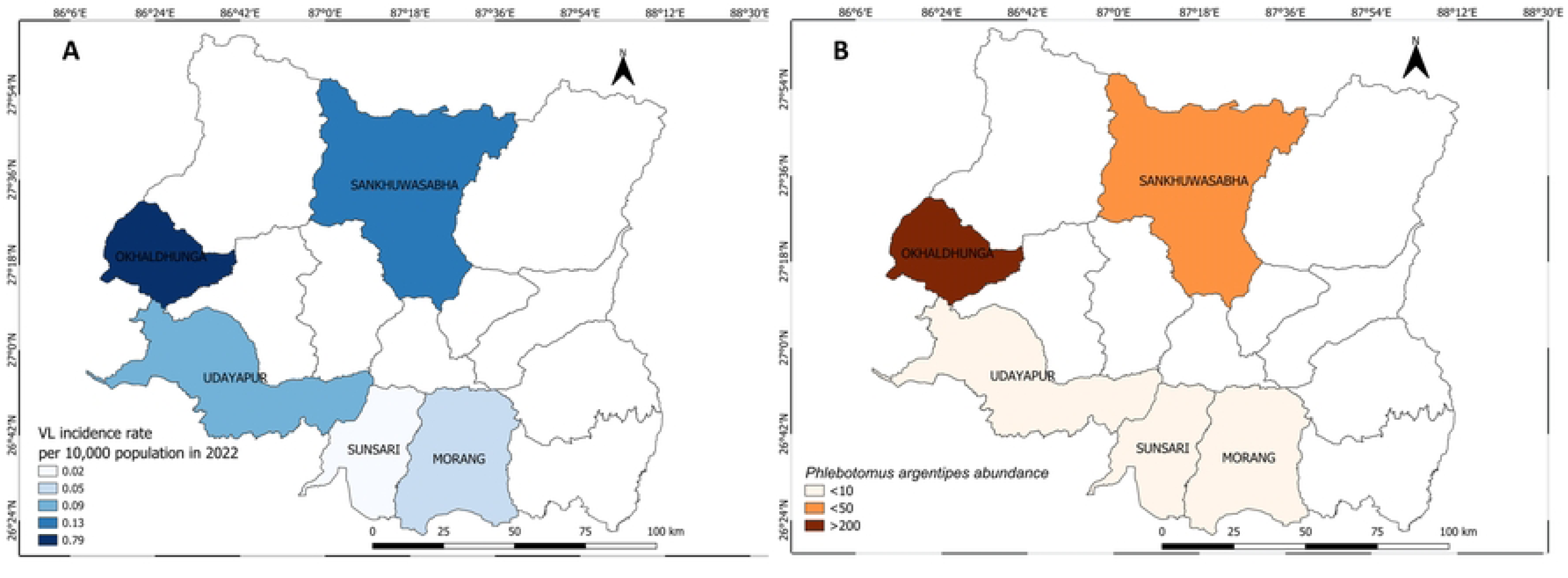
Visceral leishmaniasis incidence rate in 2022 (A) and *Ph. argentipes* abundance (B) during the survey in five study districts in Koshi province.

When fitted in the model, the disease incidence for LF was found not to be associated with the vector density possibly due to no data from two districts. However, higher *P. argentipes* density had increasing effect on VL incidence rate (IRR = 1.04, CI at 95% = 1.01 – 1.06) at district level (Table 7).

**Table 7.**
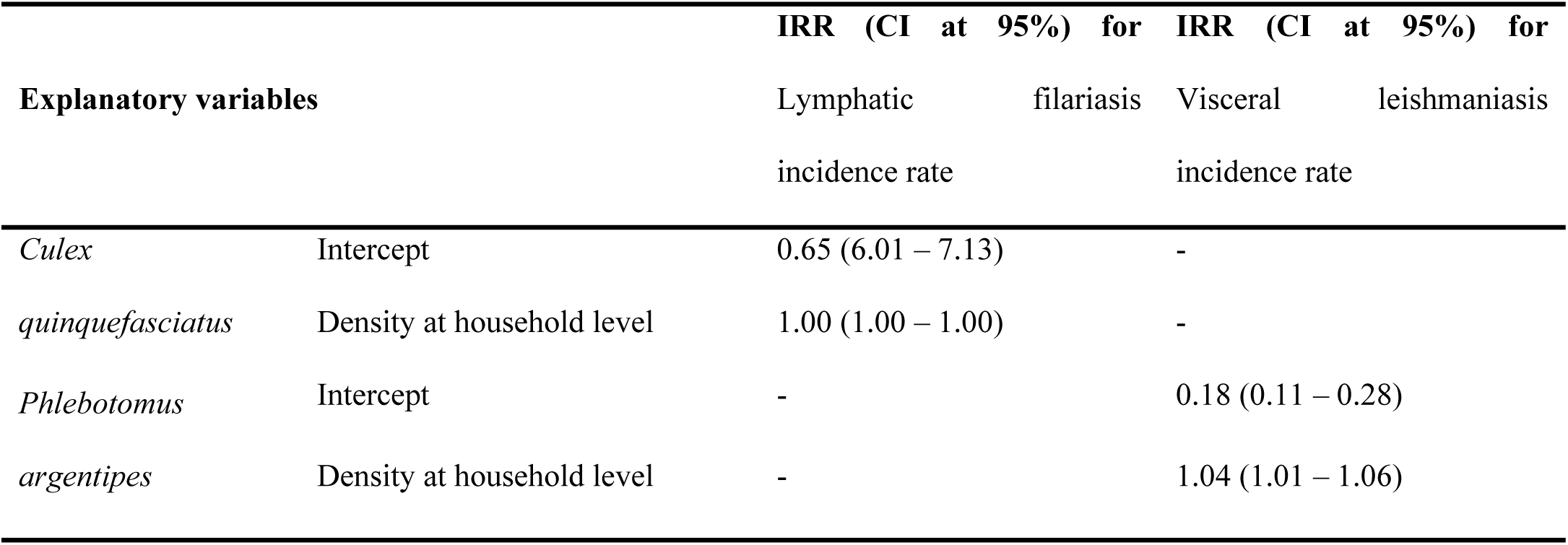
Effect of the vector density on the incidence rates of major VBDs at the district level in Koshi province.

## Discussion

Our study shows presence and abundance of the major vectors of pathogens causing malaria, VL, LF and dengue from lowland to highland districts situated at different geo-ecological and climatic regions, and distributed from in Koshi Province of Nepal. Diverse mosquito species were found in all survey districts. The dengue virus vector *Ae. aegypti* and *Ae. albopictus* were present in urban habitations of the Dharan sub-metropolitan city situated at an altitude of 300m. The *Stegomyia* indices based on the immature stages of *Aedes* spp. from the survey areas were significant enough to project intense viral transmission. Our survey documented a well-established population of *Cu. quinquefasciatus* and *Ph. argentipes* in hills and mountains, facilitated by the socio-ecological and microhabitat of these locations. In this study, we found a positive association of *Cu. quinquefasciatus* density with the mean temperature, while relative humidity had an increasing effect on the *Ph. argentipes* density. Not for all, but the vector density was positively associated with VL disease incidence in one of the survey high hill districts (Okhaldhunga).

Major malaria vector, *An. fluviatilis* was not recorded from any of the survey clusters of the five districts. Other malarial vectors, *An. maculatus* complex species; *An. willmori* and *An. pseudowillmori* were recorded from one of the high hill districts (Okhaldhunga) above 1,200m altitude. There are reports of the presence of malaria vectors from the high altitudes at 1,300m up to 2,000m in previous studies [27-29]. Abundance of *Cu. quinquefasciatus* from low altitudes to high altitudes demonstrated their resilience and adaptation to survival and establishment of their population in a wide range of geo-ecological and climatic variations in Nepal [27, 30-33]. Entomological surveillance on JE virus vector *Cu. tritaeniorhynchus* has always been meager but predictive model analysis showed the species is predominantly an Asiatic species with highly suitable environments located across Nepal, India, and China and thus supports the spread of the disease across the region [34]. In Nepal, JE is present in 63 out of 77 districts, mostly in lowlands, with some outbreaks reported in mid-hills and mountainous regions [35]. The evidence of local transmission of JE was supported by suitable ecological conditions and the presence of an abundance of vectors [27, 30, 32, 36]. The dengue virus vector, adult *Ae. aegypti* was collected in low numbers from altitudes above 300m; however, *Stegomyia* indices calculated from the immature stages of *Aedes* species indicated the threat of a potential outbreak. Previous studies conducted in Nepal have well documented the presence and abundance of *Ae. aegypti* and *Ae. albopictus* species from lowlands to high hills above 2,300m asl [4, 7, 31, 37, 38]. Another significant VBD, VL, is widely distributed in 72 endemic and endemicity doubtful districts covering all the geo-ecological and climatic regions [39], and most of these districts have a viable vector population, including high hills and mountains [40-44] (first author’s unpublished data). The findings from the current study also indicated a well-established *Ph. argentipes* density in higher altitudes. Additionally, the presence of other competent sand fly vectors has heightened the threat of VL transmission in high-altitude areas (first author’s unpublished data). A similar context of VL transmission and the evidence of the existence of potential vector species from high-altitude regions in bordering states of India [45, 46] sustains the expansion of the disease to wider geo-ecological zones.

The widespread distribution of major VBDs and their respective vectors is evident from the current epidemiological and entomological data in Nepal. Climate change over the past 40 years has been well experienced through the estimation of 0.056 °C rise in average annual maximum temperature in Nepal, with increasingly warming at higher altitudes [47]. The effect of temperature and rainfall changes strongly influence the geographical distribution of the disease and thus the VBDs [48]. An ecological time-series analysis showed 10.14% rise in VBDs hospitalizations per 1 °C rise in temperature in Nepal [49]. Another study projected 27% and 25% increase in malaria incidence with a 1 °C rise in minimum and mean temperatures, respectively [50]. Small changes in temperature and rainfall significantly influence the transmission of major VBDs. The 2018 report of the Lancet Countdown on health and climate change showed the global vectorial capacity for transmission of the dengue fever virus of *Ae. aegypti* and *Ae. albopictus* has been increased by 9.1% and 11.1% respectively in 2016 as compared to the 1950s baseline [51]. In the context of climate change, an ecological niche model in Nepal has anticipated the spread and increase in dengue virus transmission and caseload to higher altitudes [52], which is further supported by the fact of presence of vectors in these areas in this study as well.

This study has some limitations. Firstly, the entomological survey was a cross-sectional and one-time integrated vector survey in limited areas only. The results do not suffice for the generalization of diversity, distribution, and bionomics of the vectors from all the provinces in Nepal. This information is possible only if we conduct year-round surveillance of vectors in wider geographical regions. Hence, malaria vector *An. fluviatilis* was not found from any clusters of five districts during the survey. Secondly, the entomology survey was conducted in the hilly and mountainous region in the month of May and June; the peak vector-borne transmission seasons in the lowland. However, vector diversity, distribution, and bionomics can be influenced by factors like rainfall and temperature in hills and mountains; the perfect timing of vector collection in these regions remains a challenge. The prior knowledge about the seasonality of vectors in hilly and mountainous districts is crucial to set up a survey.

Sustainable integrated vector surveillance in selected sentinel sites present in various geo-ecological regions is necessary for evidence-based decision-making for the implementation of effective vector control methods. The habitat reductions and awareness to the community can be performed parallel for dengue vector surveillance. The practices of entomological survey need to be performed routinely during the non-transmission season as well. Xenomonitoring of the pathogens in the vector populations can be investigated to assess the risk of pathogen transmission in a human population living in endemic or non-endemic areas. In case of outbreaks of a particular VBDs, targeted vector surveillance should be conducted to provide evidence for prompt action on vector control and management.

## Conclusions

In conclusion, the study provides the baseline information on the diversity of vectors of major VBDs and demonstrates well-established vector populations in all geo-ecological regions with varying climatic indicators in Koshi province, Nepal. The two abundant vector species, *Cu. quinquefasciatus* and *Ph. argentipes* were indiscriminately present in lowlands to highlands (hills and mountains). In a broad spectrum, the ecological and climatic indicators are suitable for the survival, distribution and growth of vector populations. The study also examines the feasibility of integrated vector surveillance rather than a disease-specific vector surveillance for better use of the funds/resources. In addition, this study findings alert to the vector control program for regular monitoring, strengthening the existing surveillance and timely control interventions in diverse areas prone to VBD transmission.

## Acknowledgments

We would like to thank our entomology field team members: Kailash Majhi, Sashi Narayan Majhi, Satya Narayan Bhagat, Binod Uranw and Manish Karna. We are grateful to the respective district public health officials and female community health volunteers of Sunsari, Morang, Udayapur, Okhaldhunga and Sankhuwasabha for their unwavering support during the fieldwork.

## References

1. Epidemiology and Disease Control Division. National Guidelines on Integrated Vector Management: Epidemiology and Disease Control Division, Department of Health Services, Ministry of Health and Population, Government of Nepal, Teku, Kathmandu; 2020.

2. Dhimal M, Kramer IM, Phuyal P, Budhathoki SS, Hartke J, Ahrens B, et al. Climate change and its association with the expansion of vectors and vector-borne diseases in the Hindu Kush Himalayan region: A systematic synthesis of the literature. Advances in Climate Change Research. 2021;12(3):421–9. 10.1016/j.accre.2021.05.003

3. Tabachnick WJ. Challenges in predicting climate and environmental effects on vector-borne disease episystems in a changing world. J Exp Biol. 2010;213(6):946–54. 10.1242/jeb.037564

4. Dhimal M, Ahrens B, Kuch U. Climate Change and Spatiotemporal Distributions of Vector-Borne Diseases in Nepal--A Systematic Synthesis of Literature. PLoS One. 2015;10(6):e0129869. 10.1371/journal.pone.0129869 26086887: 26086887

5. Woodward A, Smith KR, Campbell-Lendrum D, Chadee DD, Honda Y, Liu Q, et al. Climate change and health: on the latest IPCC report. Lancet. 2014;383(9924):1185-9. 10.1016/s0140-6736(14)60576-6

6. Rijal KR, Adhikari B, Ghimire B, Dhungel B, Pyakurel UR, Shah P, et al. Epidemiology of dengue virus infections in Nepal, 2006–2019. Infectious Diseases of Poverty. 2021;10(1):52. 10.1186/s40249-021-00837-0

7. Dhimal M, Gautam I, Joshi HD, O’Hara RB, Ahrens B, Kuch U. Risk Factors for the Presence of Chikungunya and Dengue Vectors (Aedes aegypti and Aedes albopictus), Their Altitudinal Distribution and Climatic Determinants of Their Abundance in Central Nepal. PLOS Neglected Tropical Diseases. 2015;9(3):e0003545. 10.1371/journal.pntd.0003545

8. Shrestha UB, Gautam S, Bawa KS. Widespread Climate Change in the Himalayas and Associated Changes in Local Ecosystems. PLOS ONE. 2012;7(5):e36741. 10.1371/journal.pone.0036741

9. Ren Y-Y, Ren G, Sun X, Shrestha A, Zhan Y, Rajbhandari R, et al. Observed changes in surface air temperature and precipitation in the Hindu Kush Himalayan region over the last 100-plus years. Advances in Climate Change Research. 2017;8. 10.1016/j.accre.2017.08.001

10. Kuinkel HR. A study on spatial and temporal distribution of rainfall in Province Number 3, Nepal. Central Department of Hydrology and Meteorology: Tribhuvan University; 2019.

11. Epidemiology and Disease Control Division. National guidelines on integrated vector management. Department of Health Services, Ministry of Health and Population, Government of Nepal, Teku, Kathmandu; 2020.

12. Tyagi B, Munirathinam A, A V. A catalogue of Indian mosquitoes. International Journal of Mosquito Research. 2015;50:50–97.

13. Das Dr BP. Pictorial Key to Common Species of Culex (Culex) Mosquitoes Associated with Japanese Encephalitis Virus in India. 2013. p. 25–42.

14. Reuben R, Tewari SC, Hiriyan J, Akiyama J. Illustrated keys to species of Culex (Culex) associated with Japanese encephalitis in Southeast Asia. Mosquito Systematics. 1994;26(2).

15. Darsie RFJ, Pradhan SP, Vaidya RG. Notes on the mosquitoes of Nepal I. New country records and revised *Aedes* keys (Diptera, Culicidae). Mosquito Systematics. 1991;23(1):39–45.

16. Das Dr BP, Rajagopal R, Akiyama J. Pictorial key to the species of Indian Anophline mosquitoes. Zoology. 1990;2:131–62.

17. Darsie RFJ, Pradhan SP. The mosquitoes of Nepal: Their identification, distribution and biology. Mosquito Systematics. 1990;22(2):69–128.

18. Kalra NL, Bang YH. Manuals on Entomology in Visceral Leishmaniasis: World Health Organization, SEARO, New Delhi; 1988.

19. Lewis DJ. The Phlebotominae sandflies (Diptera: Psychodidae) of the oriental region. Bulletin of British Museum (Natural History) of Entomology. 1978;37:217–343.

20. Lewis DJ. A taxonomic review of the genus *Phlebotomus* (Diptera: Psychodidae). Bulletin of British Museum of Entomology (Natural History). 1982;45:121–209.

21. World Health Organization. Regional Office for South-East Asia. Comprehensive guideline for prevention and control of dengue and dengue haemorrhagic fever. Revised and expanded edition: WHO Regional Office for South-East Asia. New Delhi, India; 2011.

22. World Health Organization. Operational guide for assessing the productivity of Aedes aegypti breeding sites: UNICEF/UNDP/World Bank/WHO Special Programme for Research and Training in Tropical Diseases. ISBN 978-92-4-150268-9; 2011.

23. Shannon CE. A mathematical theory of communication. The Bell System Technical Journal. 1948;27(3):379–423. 10.1002/j.1538-7305.1948.tb01338.x

24. Pielou EC. The measurement of diversity in different types of biological collections. J Theor Biol. 1966;13:131–44. 10.1016/0022-5193(66)90013-0

25. Oksanen J, Simpson GL, Kindt R, Legendre P, al e. Package ‘vegan’. Community Ecology Package. Ordination methods, diversity analysis and other funtions for community and vegetation ecologists. 2022.

26. Venables WN, Ripley BD. Modern Applied Statistics with S. Fourth Edition: Springer, New York; 2002.

27. Dhimal M, Ahrens B, Kuch U. Species composition, seasonal occurrence, habitat preference and altitudinal distribution of malaria and other disease vectors in eastern Nepal. Parasites & Vectors. 2014;7(1):540. 10.1186/s13071-014-0540-4

28. Pradhan JN, Shrestha SL, Vaidya RG. Malaria transmission in high mountain valleys of west Nepal including first records of *Anopheles maculatus willmori* (James) as a third vector of malaria. J Nepal Med Assoc. 1970;8(89).

29. Pant CP, Pradhan GD, Shreshtha SL. Distribution of Anophelines in relation to altitude in Nepal. World Health Organization, Geneva: WHO/Mal/343; 1962.

30. Maharjan M, Pant S, Pant D. Distribution of Mosquito species in Kathmandu, Rupandehi, Kapilbastu and Morang Districts of Nepal. 2014;20:1.

31. Dhimal M, Gautam I, Kreß A, Müller R, Kuch U. Spatio-Temporal Distribution of Dengue and Lymphatic Filariasis Vectors along an Altitudinal Transect in Central Nepal. PLoS Negl Trop Dis. 2014;8(7):e3035. 10.1371/journal.pntd.0003035

32. Shrestha M, Gautam I, Gupta R. Study on Culex mosquitoes of Bhelukhel, Bode and Tathali of Bhaktapur district, Nepal. J Nat Hist Mus 2014;28:118–26.

33. Byanju R, Gautam I, Aryal M, Kc A, Shrestha HN, Dhimal M. Adult Density of Culex quinquefasciatus Say, Filarial Vector in Thapa Gaun, Jhaukhel and Lama Tole, Nagarkot VDC, Bhaktapur District. Nepal Journal of Science and Technology. 2013;14(1):185–94. 10.3126/njst.v14i1.8939

34. Longbottom J, Browne AJ, Pigott DM, Sinka ME, Golding N, Hay SI, et al. Mapping the spatial distribution of the Japanese encephalitis vector, Culex tritaeniorhynchus Giles, 1901 (Diptera: Culicidae) within areas of Japanese encephalitis risk. Parasites & Vectors. 2017;10(1):148. 10.1186/s13071-017-2086-8

35. Kumar Pant D, Tenzin T, Chand R, Kumar Sharma B, Raj Bist P. Spatio-temporal epidemiology of Japanese encephalitis in Nepal, 2007-2015. PLoS One. 2017;12(7):e0180591. 10.1371/journal.pone.0180591

36. Tamrakar AS. Seasonal distribution of *Culex tritaeniorhynchus* Giles (Diptera: Culicidae), the vector of Japanese encephalitis in Kathmandu valley. Kathmandu: Nepal Health Research Council eLibrary; 2009.

37. Oli B, Sharma M, Shrestha P, Dhimal M, Gautam I. Breeding Habitat Preference of Aedes aegypti (Linnaeus, 1762) and Aedes albopictus (Skuse, 1895) along an Altitudinal Gradient in Mid-Western Nepal. Journal of Institute of Science and Technology. 2024;29:83–91. 10.3126/jist.v29i1.64658

38. Gautam I, Dhimal M, Shrestha S, Tamrakar A. First Record of Aedes aegypti (L.) Vector of Dengue Virus from Kathmandu, Nepal. J Nat Hist Mus. 2009;24:156–64. 10.3126/jnhm.v24i1.2298

39. Jain S, Madjou S, Agua JFV, Maia-Elkhoury AN, Valadas S, Warusavithana S, et al. Global leishmaniasis surveillance updates 2023: 3 years of the NTD road map. World Health Organization; 2024 8 November 2024. Report No.: 45.

40. Uranw S, Bhattarai NR, Cloots K, Roy L, Rai K, Kiran U, et al. Visceral leishmaniasis in the hills of western Nepal: A transmission assessment. PLoS One. 2024;19(4):e0289578. 10.1371/journal.pone.0289578

41. Roy L, Cloots K, Uranw S, Rai K, Bhattarai NR, Smekens T, et al. The ongoing risk of Leishmania donovani transmission in eastern Nepal: an entomological investigation during the elimination era. Parasites & Vectors. 2023;16(1):404. 10.1186/s13071-023-05986-9

42. Joshi AB, Banjara MR, Das ML, Ghale P, Pant KR, Pyakurel UR, et al. Epidemiological, Serological, and Entomological Investigation of New Visceral Leishmaniasis Foci in Nepal. Am J Trop Med Hyg. 2023. 10.4269/ajtmh.23-0373

43. Ostyn B, Uranw S, Bhattarai NR, Das ML, Rai K, Tersago K, et al. Transmission of Leishmania donovani in the Hills of Eastern Nepal, an Outbreak Investigation in Okhaldhunga and Bhojpur Districts. PLoS Negl Trop Dis. 2015;9(8):e0003966. 10.1371/journal.pntd.0003966

44. Uranw S, Hasker E, Roy L, Meheus F, Das ML, Bhattarai NR, et al. An outbreak investigation of visceral leishmaniasis among residents of Dharan town, eastern Nepal, evidence for urban transmission of Leishmania donovani. BMC Infect Dis. 2013;13:21. 10.1186/1471-2334-13-21

45. Lata S, Kumar G, Ojha VP, Dhiman RC. Detection of Leishmania donovani in Wild-Caught Phlebotomine Sand Flies in Endemic Focus of Leishmaniasis in Himachal Pradesh, India. J Med Entomol. 2022;59(2):719–24. 10.1093/jme/tjab202

46. Sharma NL, Mahajan VK, Ranjan N, Verma GK, Negi AK, Mehta KI. The sandflies of the Satluj river valley, Himachal Pradesh (India): some possible vectors of the parasite causing human cutaneous and visceral leishmaniases in this endemic focus. J Vector Borne Dis. 2009;46(2):136–40.

47. Pandey BD, Costello A. The dengue epidemic and climate change in Nepal. The Lancet. 2019;394(10215):2150–1. 10.1016/S0140-6736(19)32689-3

48. Fouque F, Reeder JC. Impact of past and on-going changes on climate and weather on vector-borne diseases transmission: a look at the evidence. Infectious Diseases of Poverty. 2019;8(1):51. 10.1186/s40249-019-0565-1

49. Shrestha SL, Shrestha IL, Shrestha N, Joshi RD. Statistical Modeling of Health Effects on Climate-Sensitive Variables and Assessment of Environmental Burden of Diseases Attributable to Climate Change in Nepal. Environmental Modeling & Assessment. 2017;22(5):459–72. 10.1007/s10666-017-9547-5

50. Dhimal M, O’Hara RB, Karki R, Thakur GD, Kuch U, Ahrens B. Spatio-temporal distribution of malaria and its association with climatic factors and vector-control interventions in two high-risk districts of Nepal. Malar J. 2014;13(1). 10.1186/1475-2875-13-457

51. Watts N, Amann M, Arnell N, Ayeb-Karlsson S, Belesova K, Berry H, et al. The 2018 report of the Lancet Countdown on health and climate change: shaping the health of nations for centuries to come. Lancet. 2018;392(10163):2479-514. 10.1016/s0140-6736(18)32594-7

52. Acharya BK, Cao C, Xu M, Khanal L, Naeem S, Pandit S. Present and Future of Dengue Fever in Nepal: Mapping Climatic Suitability by Ecological Niche Model. International Journal of Environmental Research and Public Health. 2018;15(2):187.

